# Intrinsic properties of spinal motoneurons degrade ankle torque control in humans

**DOI:** 10.1101/2023.10.23.563670

**Authors:** James. A. Beauchamp, Gregory E. P. Pearcey, Obaid U. Khurram, Francesco Negro, Julius P.A. Dewald, CJ. Heckman

## Abstract

Motoneurons are the final common pathway for all motor commands and possess intrinsic electrical properties that must be tuned to control muscle across the full range of motor behaviors. Neuromodulatory input from the brainstem is likely essential for adapting motoneuron properties to match this diversity of motor tasks. A primary mechanism of this adaptation, control of dendritic persistent inward currents (PICs) in motoneurons by brainstem monoaminergic systems, generates both amplification and prolongation of synaptic inputs. While essential, there is an inherent tension between this amplification and prolongation. Although amplification by PICs allows for quick recruitment and acceleration of motoneuron discharge during discrete motor tasks, PICs must be deactivated to de-recruit motoneurons upon movement cessation. In contrast, during stabilizing or postural tasks, PIC-induced prolongation of synaptic inputs is likely critical for sustained motoneuron discharge. Here, we designed two motor tasks that show PIC amplification and prolongation may conflict and generate errors that degrade the precision of motor output in humans. This included a paradigm comprised of a discrete motor task superimposed atop a stabilizing task and a paradigm with muscle length-induced changes to the balance of excitatory and inhibitory inputs available for controlling PICs. We show that prolongation from PICs introduces deficits in ankle torque control and that these deficits are further degraded at shorter muscle lengths when PIC prolongation is greatest. These results highlight the necessity for inhibitory control of PICs and showcase issues that are introduced when inhibitory control is perturbed or constrained. Our findings suggest that, like sensory systems, errors are inherent in motor systems. These errors are not due to problems in the perception of movement-related sensory input but are embedded in the final stage of motor output.

## Introduction

Alpha motoneurons are the final common output of the central nervous system and transduce all motor commands (i.e., synaptic inputs) into discharge patterns for the generation of force (Sherrington, 1907, Heckman and Enoka, 2012). Thus, it is reasonable to assume that the intrinsic electrical properties that govern this transduction are adapted over a wide range to facilitate the large assortment of motor behaviors observed in humans (Khurram et al., 2022, Johnson et al., 2017). However, insight into how motoneurons effectively implement motor commands has been complicated by the discovery that their intrinsic electrical properties are profoundly altered by brainstem-originating neuromodulatory inputs (e.g., norepinephrine, serotonin) (Schwindt and Crill, 1977, Schwindt and Crill, 1980, Hounsgaard et al., 1984, Lee and Heckman, 1998, Lee and Heckman, 2000).

Brainstem axons that release the monoamines serotonin (5HT) or norepinephrine (NE) form dense, monosynaptic projections to the spinal cord that control the excitability of motoneurons (Holstege and Kuypers, 1987, Bowker et al., 1982, Alvarez et al., 1998). In adult motoneurons, 5HT and NE facilitate voltage-sensitive sodium (Na) and calcium (Ca) channels that mediate persistent inward currents (PICs) (Udina et al., 2010, Lee and Heckman, 1999, Hounsgaard et al., 1988, Hounsgaard and Kiehn, 1985). PICs provide an additional source of depolarizing (i.e., excitatory) current that both amplifies and prolongs synaptic inputs to motoneurons by upwards of 3-5 fold, imparting distinct characteristics to their discharge patterns (Binder et al., 2020, Lee and Heckman, 1996, Lee and Heckman, 2000). These distinct characteristics are clearly manifest in human motor unit discharge patterns and are likely fundamental for normal motor output (Khurram et al., 2022, Johnson et al., 2017, Heckman and Enoka, 2012).

Brainstem neuromodulation of PICs by monoamines, and the resulting amplification and prolongation of excitatory synaptic inputs, have multiple important functions. Amplification from PICs boosts the modest synaptic currents generated by individual excitatory input systems so that each can strongly affect motoneuron discharge (Binder and Powers, 2001, Heckman et al., 2008b, Cushing et al., 2005). Likewise, in parallel, the prolongation of discharge provided by PICs is likely essential for postural tasks and underlies the self-sustained discharge of motoneurons that is observed following brief excitatory inputs in intracellular recordings (i.e., bistable behavior) (Crone et al., 1988, Hounsgaard et al., 1988, Heckman et al., 2008b). Consequently, neuromodulation of PICs provides the basis for gain control of motor output, in which variations in monoamines adapt motoneuron excitability to match the widely varying force demands of the normal motor repertoire (Johnson and Heckman, 2014, Naufel et al., 2019, Wei et al., 2014)

While essential, there is an inherent tension between the amplification and prolongation of excitatory inputs provided by PICs. Though amplification by PICs allows for quick recruitment and acceleration of motoneuron discharge, these PICs must be deactivated to de-recruit motoneurons and avoid sustained activation upon movement cessation. In contrast, continued activation of PICs (i.e., prolongation) is likely necessary for the sustained contractions inherent to stabilization and postural tasks. These contrasting task-dependent demands necessitate a precise control mechanism to adjust their magnitude and duration. While monoaminergic drive adjusts the magnitude of PICs, these effects lack the specificity necessary to independently adjust PIC amplification and prolongation. Thus, contrasting task demands are likely accomplished by precise inhibitory control embedded within motor commands (Heckman and Enoka, 2012, Hyngstrom et al., 2007, Heckman et al., 2008a, Johnson et al., 2012, Bui et al., 2008a, Kuo et al., 2003). We, therefore, reasoned that much like sensory systems, where perceptual errors can occur when presented with conflicting inputs, tasks that push the capacity of inhibitory inputs to control PIC behaviors would result in aberrant discharge patterns and poorly controlled forces.

We designed two isometric contraction paradigms that were likely to pose this type of challenge. In the first paradigm, we superimposed a discrete motor task and a prolonged low-effort stabilization task. In this superimposition task (colloquially termed sombrero), the inhibitory input necessary to deactivate the PICs of higher threshold motoneurons recruited during the discrete task is compromised because of the sustained excitatory input required for the stabilizing task. We, therefore, hypothesized that PIC prolongation of higher threshold motor units, otherwise not required for this low-level output, would be excessive after the return to stabilization and result in highly variable torque control. In the second paradigm, we modulated the balance of excitatory and inhibitory input available for controlling PICs by changing the length of agonist-antagonist muscle pairs prior to isometric motor tasks. Studies in humans have highlighted an increase in spinal excitability as a muscle is shortened, likely yielding a relative reduction in net inhibitory input to the motor pool and hindering PIC deactivation (Hwang, 2002, Frigon et al., 2007, Dutt-Mazumder et al., 2020, Patikas et al., 2004). Thus, we hypothesized that motoneuron outputs would suffer from excessive PIC prolongation at short agonist muscle lengths, again resulting in poor torque control.

Our results strongly supported our hypothesis for the first paradigm and revealed excessive PIC prolongation and poor torque control following cessation of the discrete task and return to the stabilizing task. Similarly, in the second paradigm, we found greater PIC prolongation at short agonist lengths for both ankle dorsiflexors and plantar flexors. Combining these two paradigms further supported this finding and highlighted greater PIC prolongation and worsened torque control during the superimposition task when a muscle is in a shortened position.

Though the monoaminergic dependence of PICs allows for brainstem control of motoneuron excitability, the inherent characteristics of PICs necessitate task-dependent inhibitory control. Here we have highlighted this necessity and shown two instances where conflicts and/or constraints to inhibitory control of PICs introduce quantifiable difficulties in human torque control. These findings have many implications with relevance to clinical conditions where pathological shifts in monoamines are theorized (e.g., stroke, SCI) and may potentially shed light on contributing factors to muscle cramps (Li et al., 2019, Murray et al., 2010, McPherson et al., 2018b, Beauchamp et al., 2022a). It is often appreciated that the prevalence of muscle cramps increases at shortened muscle lengths, and PICs have even been theorized as a causative factor, though an empirical link between PICs and muscle cramps has remained unestablished (Layzer and Rowland, 1971, Schwellnus, 2009, Baldissera et al., 1991, Minetto et al., 2013). Here we show that prolongation from PICs is greatest at shorter muscle lengths and may impede volitional relaxation of muscle, forcing sustained muscle contraction as is observed in muscle cramps.

## Methods

### Overview

The primary goal of this work was to highlight instances where the intrinsic properties of spinal motoneurons (i.e., persistent inward currents; PICs) introduce difficulties in human motor control, due to their need for task-dependent inhibitory control. To accomplish this, we designed two experimental paradigms that introduced potential constraints and conflicts to inhibitory control of PICs. We then used high-density surface electromyography (HDsEMG) to estimate human motor unit (MU) behavior and PIC characteristics during these paradigms and observed the subsequent behavioral ramifications. The first paradigm is presented as experiment one, the second paradigm as experiment two, and a combination of both paradigms as experiment three.

### Experimental Setup

For each paradigm, similar configurations were employed. In all sessions, participants were seated in a Biodex chair, and their left foot was securely attached to a footplate fixed onto a Systems 4 Dynamometer (Biodex Medical Systems, Shirley, NY) with thigh and shoulder straps employed to minimize movement. Throughout each session, the center of rotation of a participant’s ankle joint was aligned with the dynamometer’s axis of rotation and their left knee and hips were maintained at 20° and 80° of flexion, respectively. In experiment one, a participant’s ankle was placed at 90° and was adjusted ±20° in experiments two and three, as detailed in the protocol. Target isometric contractions and visual feedback (i.e., ankle dorsiflexion or plantarflexion torque) were provided on a flat monitor via a custom Matlab interface (MATLAB (R2020b), The Mathworks Inc., Natick, MA). Torque about the ankle was filtered with a 125 ms moving average window before being provided as visual feedback to the participant. For subsequent analysis, raw torque signals were digitized (2048 Hz) using a 16-bit analog-to-digital converter (Quattrocento, OT Bioelettronica, Turin, IT) and lowpass filtered (50 Hz; fifth-order Butterworth filter).

### Experimental Protocol

To streamline data collection, the paradigms presented as experiments one, two, and three were conducted on the same day in random order for each participant. To normalize subsequent efforts, each experimental session began with determining a participant’s maximal isometric torque generating capacity for both dorsiflexion and plantarflexion. This was done independently for each of the three ankle angles to be tested, with the ankle positioned at 70°, 90°, or 110°. For each ankle angle and torque direction, a minimum of two maximal contractions were performed, with at least one minute of rest separating contractions. Maximal contractions were repeated until the peak torque within the last contraction deviated by less than 10% of the previous two contractions. We then used the maximum voluntary torque (MVT) achieved during these efforts to normalize all subsequent contractions. Prior to each paradigm, a minimum of six practice trials were performed, to ensure that participants understood and could adequately perform the desired contraction paradigms. Practice trials were performed until an individual could match the desired torque trajectories with less than ±5% error for more than half the trial, with real-time visual feedback of an individual’s performed torque and desired torque provided. Both paradigms were carried out in the dorsiflexion and plantarflexion direction with HDsEMG recorded from surface electrode arrays atop the skin overlying the tibialis anterior (TA) and medial gastrocnemius (MG) muscle bellies.

*Experiment 1:* The data presented in experiment one consists entirely of the first paradigm. In the first paradigm, we constrained the inhibitory input necessary to deactivate PICs by requiring individuals to maintain excitatory drive to their motor pools. In short, individuals produced a linear increase and decrease in isometric ankle torque (i.e., ramp contraction) while performing a low-effort stabilizing isometric ankle torque task. Practically, for each trial, individuals were first asked to generate isometric dorsiflexion or plantarflexion ankle torque to 10% MVT and maintain this effort for 10 s, as precisely and accurately as possible (referred to as *plateau one*). Following 10 s, individuals were then asked to linearly increase and decrease their effort by approximately 3% MVT/s to 30% MVT and back to 10% MVT (referred to as *ramp*). Following their return to 10% MVT, individuals were again asked to hold 10% MVT for 10 s, as precisely and accurately as possible (referred to as *plateau two*). Colloquially, due to the shape of this desired torque trajectory and their resemblance to a 2-D sombrero function and/or hat, these have been referred to as sombrero contractions in conference proceedings and communications with colleagues. Control trials, where individuals were required to maintain 10% MVT for the duration of the contraction instead of the linear increase and decrease to 30% MVT (i.e., sustained hold), were randomly interspersed. A minimum of four sustained hold and sombrero contractions were performed, two each for plantarflexion or dorsiflexion. Throughout all trials, antagonist EMG was monitored, and participants were coached to use only agonist muscles for the intended dorsiflexion or plantarflexion contraction.

During sombrero contractions, the additional excitatory synaptic input necessary to recruit higher threshold MUs and perform the linear ramp contraction also activates the PICs in these recruited units. While PICs are indeed necessary for repetitive discharge, due to the slow inactivation of the dendritic channels that mediate them, PICs keep a motoneuron suprathreshold and prolong its discharge to lower levels of excitatory current than originally necessary for its recruitment. To attenuate this prolongation, inhibitory inputs could be used to shunt the excitatory dendritic current from PICs. During a traditional ramp contraction (i.e., linear increase and decrease in effort) this would be without problem, as individuals could simply increase inhibitory input or cease all excitatory input to deactivate PICs and de-recruit MUs. By requiring that individuals slowly relax to a maintained low-level effort (i.e., *plateau two*; 10% MVT hold), we are requiring that individuals maintain a base level of excitatory drive to the motor pool. This excitatory drive is by nature in conflict with the inhibitory inputs necessary to inactivate PICs in the MUs that were recruited for the ramp contraction. Thus, the PICs in these MUs force sustained discharge well into the second 10% MVT hold (*plateau two*). Given that these higher threshold MUs were originally recruited above 10%, many higher threshold MUs are now actively discharging during the second 10% MVT plateau and individuals must alter their control strategy.

*Experiment 2:* The data presented in experiment two is comprised of the second paradigm, in which we modulated the balance of excitatory and inhibitory input to the motor pool, presumably altering the degree of inhibition available for PIC deactivation. Prior studies have shown an increase in spinal excitability as a muscle is shortened, with increases in Hmax/Mmax ratio (Hwang, 2002, Frigon et al., 2007, Dutt-Mazumder et al., 2020). Increasing spinal excitability likely yields a relative reduction of inhibitory input and/or disinhibition to motoneurons, which would reduce inhibitory constraints on PICs. Prior work from our group has highlighted the unique sensitivity of PICs to inhibitory inputs, due to their dendritic location, and even shown ankle joint angle in the decerebrate cat to modulate PIC magnitude (Hyngstrom et al., 2007, Heckman et al., 2008a, Kuo et al., 2003). Thus, we reasoned that changing the length of agonist muscles should either constrain (long muscle, reduced spinal excitability) or facilitate (short muscle, increased spinal excitability) PICs. To accomplish the desired change in agonist-antagonist muscle pairs we modulated the angle of the ankle prior to various isometric dorsiflexion and plantarflexion tasks. Individuals were asked to produce linear isometric dorsiflexion and plantarflexion ramp contractions to 30% MVT with a rise and decay speed of 3% MVT/s. This was done in random order for three separate ankle angles (70°, 90°, and 110°). At 70°, the TA is in a shortened position while the MG and SOL are relatively lengthened. Whereas, at 110°, the MG and SOL are in a shortened position while the TA is in a lengthened position. For each ankle angle, a minimum of four ramp contractions were performed, two each for plantarflexion or dorsiflexion. For this experiment, we performed linear isometric ramp contractions, as these contractions are widely used in the field (Pearcey et al., 2022, Orssatto et al., 2023, Jenz et al., 2023, Lapole et al., 2023) and have well validated metrics for estimating PICs in humans (Gorassini et al., 2002, Afsharipour et al., 2020, Beauchamp et al., 2023, Chardon et al., 2023), and in specific prolongation of discharge from PICs. All trials were adjusted for the maximum voluntary torque generated in the exact configuration to be tested. For example, ramp contractions conducted at 70° peaked at 30% of the maximum voluntary torque generated in either dorsiflexion or plantarflexion while the ankle is at 70°. Throughout all trials, antagonist EMG was monitored and participants were coached to use only agonist muscles for the intended dorsiflexion or plantarflexion contraction.

*Experiment 3:* The data presented in experiment three is a combination of both experimental paradigms. Given that changes in the balance of excitation-inhibition were expected from changes in agonist muscle length, we speculated that we could leverage the constraint and/or facilitation of PICs achieved at various ankle joint angles to adjust the PIC prolongation observed in the first experiment. That is, given that excessive PIC prolongation yields sustained discharge of higher threshold MUs and difficulties in force control in the sombrero task, we reasoned that increasing spinal excitability would further hinder PIC deactivation and degrade torque control. In contrast, reducing spinal excitability should assist PIC deactivation and improve torque control. To probe this question, we asked individuals to perform sombrero tasks or sustained holds at the two extreme ankle angles of experiment two (i.e., 70° and 90°). A minimum of sixteen contractions were performed: two replicates for plantarflexion or dorsiflexion at both ankle angles for the sombrero contractions and sustained holds. As before, the desired torque trajectories supplied to participants were normalized to the maximum voluntary dorsiflexion or plantarflexion torque for a given ankle angle.

### Motor Unit Decomposition

High-Density Surface EMG (HDsEMG) was collected with 64 channel electrode grids (GR08MM1305, OT Bioelettronica, Turin, IT) placed atop the skin overlying the TA and MG muscle bellies with adhesive foam and conductive paste. Prior to electrode placement, the muscles of interest were identified by experienced investigators and the skin overlying the muscle was shaved, abraded with abrasive paste, and cleaned with isopropyl alcohol. Two Ag/AgCl ground electrodes were placed bilaterally on the right and left patella and a moist band electrode was placed around the right ankle. HDsEMG was acquired with differential amplification (150 x), bandpass filtered (10-900 Hz), and digitized (2048 Hz) using a 16-bit analog-to-digital converter (Quattrocento, OT Bioelettronica, Turin, IT).

Following collection, each channel of surface EMG was visually inspected to remove channels with substantial artifacts, noise, or saturation of the A/D board. The remaining EMG channels were decomposed into individual MU spike trains using convolutive blind source separation and successive sparse deflation improvements (Negro et al., 2016, Martinez-Valdes et al., 2017). The silhouette threshold for decomposition was set to 0.87. To improve decomposition accuracy, automatic decomposition results were augmented by iteratively re-estimating the spike train and correcting for missed spikes or substantial deviations in the discharge profile (Martinez-Valdes and Negro, 2023).

### Participant Population

In total, analysis for experiment one included 1327 MUs for the TA and 1157 MUs for the MG. For experiment two, this included 2035 total MUs the TA and 1976 MUs for the MG. Likewise, experiment three included 1959 MUs for the TA and 2128 MUs for the MG across both ankle angles tested. Data was collected from twelve healthy volunteers (26.5 ± 3.0 years; 2F, 10M) with no known history of cardiovascular, metabolic, or neuromuscular impairment for each experiment. All individuals provided written and informed consent prior to participation, in accordance with Northwestern University Institutional Review Board (STU00202964).

### Data Analysis

Following decomposition, binary MU spike trains were used to generate discrete estimates of instantaneous discharge rate. For each MU, instantaneous estimates were then smoothed with support vector regression to create continuous estimates, as previously described (Beauchamp et al., 2022b). In brief, the reciprocal of the inter-spike intervals (ISI), or the time between consecutive spikes, were used to train a support vector regression (SVR) model with L1 soft-margin minimization to predict instantaneous discharge rate as a function of the corresponding time instances for each MU (MATLAB (R2020b), The Mathworks Inc., Natick, MA). Smooth and continuous estimates of discharge rate were then generated with this SVR model along a time vector from MU recruitment to derecruitment sampled at 2048 Hz (MATLAB (R2020b), The Mathworks Inc., Natick, MA). Hyperparameters were chosen in accordance with those previously suggested (Beauchamp et al., 2022b, Beauchamp et al., 2023).

### Estimates of PIC-induced prolongation of discharge

For the sombrero contractions, estimates of PIC prolongation were facilitated by separating MUs into three cohorts based upon the torque at which they were recruited during the contraction. This included brim MUs that were recruited at the onset of the stabilization task in *plateau one*, button MUs that were recruited and derecruited in the center ramp region, and cap MUs which were recruited in the center ramp but sustained discharge into the second plateau. The duration of sustained discharge (i.e., prolongation) from PICs was estimated as the duration of time that a cap MU maintained discharge (i.e., ISI < 1 s) after its theoretically expected point of derecruitment. These values are calculated as the duration of time (s) that a MU continues to discharge past the instance that it should have been derecruited if its recruitment and derecruited torque were identical. That is, if a MU was recruited at 15% MVT on the ascending portion of the center ramp, its sustained discharge would be represented by the time that it continued to fire past 15% MVT on the descending portion of the ramp. Furthermore, the proportion of cap MUs that sustained discharge was quantified per trial as the total number of cap MUs that sustained discharger greater than 2 s, divided by the total number of MUs recruited in the ramp contraction (i.e., cap + brim MUs). MU recruitment and derecruitment was estimated as the time (and torque) at which the decomposition predicted the first and last instantaneous MU discharge, respectively.

In experiment two, we employed a collection of methods to quantify PIC prolongation. These metrics are detailed below and include a paired MU analysis technique (ΔF), the duration of discharge that a MU exhibits on the descending phase of the ramp contraction, and the torque at which MUs were derecruited. An additional battery of metrics was conducted and can be found in the supplementary information for the interested reader.

#### Paired Motor Unit Analysis

delta frequency (ΔF) is a commonly employed and well characterized metric used to estimate the magnitude of PICs and represents the hysteresis of a higher threshold MU with respect to the discharge rate of a lower threshold unit. ΔF for a given MU (test unit) is quantified as the change in discharge rate of a lower threshold MU (reporter unit) between the recruitment and derecruitment instance of the test unit. To account for the possible pairing of a test unit with multiple lower threshold reporter units, we represented ΔF for a given test unit as the average change in discharge rate across all reporter unit pairs. To ensure the validity of ΔF estimates, we employed multiple exclusion criteria for test-reporter unit pairs; to ensure that MU pairs likely received a common synaptic drive, we only included test unit-reporter unit pairs with rate-rate correlations of r^2^ > 0.7 (Gorassini et al., 2002, Udina et al., 2010, Wilson et al., 2015); to ensure full activation of the PIC in the reporter unit, we excluded any pairs with recruitment time differences <1 s (Powers et al., 2008, Bennett et al., 2001, Hassan et al., 2020); and to avoid saturated reporter units, we excluded test unit-reporter unit pairs in which the reporter unit discharge range was < 0.5 pps while the test unit was active (Stephenson and Maluf, 2011).

#### Descending discharge and derecruitment

To provide insight into the behavior of MUs on the descending phase of discharge, we extracted two key metrics. This included the total duration of time that a MU exhibited sustained discharge following peak torque. Quantitatively, this was calculated as the time in seconds from the occurrence of peak torque for a given trial to the time of derecruitment for each MU. To further characterize the descending phase of MU discharge we also quantified the torque at which each MU was derecruited and represent this as a percentage of maximum voluntary torque.

#### Supplemental metrics

Additional metrics not highlighted in the primary results for the second experiment, but of potential interest, can be found in the supplement information and include torque at MU recruitment, torque at MU derecruitment, the duration of MU discharge on the ascending phase of the ramp, the duration of discharge on the descending phase of the ramp, the difference in ascending and descending duration as a function of the total duration of MU discharge (i.e. ADR: ascending-descending ratio) (Afsharipour et al., 2020, Hassan et al., 2021), and the discharge rate at MU recruitment, peak, and derecruitment.

### MU Matching

To observe the change in metrics employed in the second experiment for presumably the same MU across each muscle length, we identified repeated observations of the same MU at each length. Identifying the same MU at each joint angle (muscle length) allows for estimating PIC prolongation whilst removing the potential bias that may be introduced by decomposing and/or recruiting varying populations of MUs at each length. To do this, we estimated MU action potential waveforms (MUAPs) with spike triggered averaging and computed a 2-D cross correlation between the spatial representation of the MUAPs between ramp contraction trials (Martinez-Valdes et al., 2017, Del Vecchio et al., 2019). This was repeated across all MUs in successive contractions at each muscle length, with normalized correlation values between MU pairs greater than 0.8 deemed a matched unit. Matched unit pairs across trials were given a single unique MU identifier. We then collected the values for each of the proposed metrics for matched MUs to estimate the change in MU discharge introduced by changes in muscle length. Outcome metrics for only MUs matched at a minimum of two muscle lengths can be observed in the supplementary information. This matched data follows the same trends of the unmatched MUs displayed in the results.

### MU Coherence

To observe potential alterations in common drive to the motor pool, we performed a coherence analysis for MUs discharging in each plateau of the sombrero task. For each sombrero trial, this included separating the trial into two segments indicating the first and second plateau and collecting the spike trains of MUs actively discharging in either of these regions. Separately for each plateau, this collection of MUs was then randomly sampled, creating two groups of four MUs. The detrended sum of all spike trains for each group of MUs was then computed and the cross-spectra and auto-spectra were quantified (*no overlap, NFFT = 10*sampling rate, window length = 1 s*). This was repeated for 100 iterations, with common drive indicated as the product of the summed cross spectra over the product of the summed auto spectra. Similarly, we performed this analysis between joint torque and MU discharge, using the same approach but for each of the 100 iterations quantifying the cross and auto-spectra between four randomly sampled MUs and the corresponding joint torque. For both approaches, the spectra were z-transformed as depicted in the equation below (Rosenberg et al., 1989, Baker et al., 2003), where L indicates the number of disjoint segments, and the bias represents the average transformed coherence from 500 – 750 Hz.

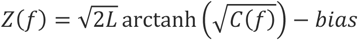

These transformed values were then segmented into bandwidths (delta [0.5–5 Hz], theta [5–10 Hz], alpha [10–15 Hz], beta [15–25 Hz]), with the average magnitude of the low frequency delta component reported.

### Statistical Analysis

To observe task performance and MU characteristics within the sombrero and hold contractions, each trail was segmented into two regions indicating either *plateau one* or *plateau two*. These regions of times were matched between the sombrero and hold contractions. To compare changes between plateaus, for the sombreros and holds independently, we quantified the average discharge rate of MUs and coefficient of variation in torque for each plateau and fit each of these metrics with a linear mixed model that contained fixed effects of a muscle (MG, TA) and plateau region (1, 2), with a random effect of participant and covariate of trial number. To appreciate the magnitude of change between plateaus, we computed estimated marginal mean differences between plateau factor levels. To compare differences between the sombrero and hold contractions, we quantified the change in each metric from *plateau one* to *two* (Δ’s) for each trial and fit a linear mixed model to these differences, comprised of fixed effects of muscle (MG, TA), contraction type (sombrero, hold), and their interaction with a random effect of participant and covariate of trial number. We then computed estimated marginal means for these Δ values and quantified the effect size (Cohen’s *d*) for the difference in these values between the sombrero and hold contractions.

To observe changes in the prolongation of MU discharge as a function of muscle length, we fit a linear mixed model to the battery of aforementioned metrics with fixed effects of muscle (MG, TA), length (long, mid, short), and their interaction. For the unmatched population data, we employed a random effect of participant and covariate of trial number. For the matched data, we employed a random effect of MUid (a unique identifier of independent MUs). We then computed estimated marginal means for each metric and quantified the effect size (Cohen’s *d*) for the difference in these values between the long and short lengths.

To observe changes in cap MU behavior as a function of length, we quantified the sustained duration for each cap MU and fit a linear mixed model to these values comprised of fixed effects of muscle (MG, TA), length (long, short), and their interaction with random effects of participant and covariate of trail number. Similarly, for each trial we quantified the number of cap MUs that sustained discharge >2s as a function of the button MUs, the total number of cap MUs, and their average discharge rate and fit a linear mixed model to these values comprised of fixed effects of muscle (MG, TA), length (long, short), and their interaction with random effects of participant. To observe task performance and MU characteristics within the sombrero and hold contractions, we conducted a similar protocol to experiment one. Explicitly, we quantified the change in torque and average MU discharge rate between *plateau one* and *two* for each trial and fit with a linear mixed model with fixed effects of contraction type (sombrero, hold), muscle (MG, TA), length (long, mid, short) and their interaction with a random effect of participant and covariate of trail number. We then computed estimated marginal means for each metric and quantified the effect size (Cohen’s *d*) for the difference in these values between the long and short lengths and between contraction types.

All statistical analysis was performed with R (R Core Team, 2021). Mixed model analysis was achieved via the lme4 (Bates et al., 2015) package and p-values were obtained by likelihood ratio tests of the full model with the effect in question against the model without the effect in question. For main effects, this included their subsequent interaction terms. To ensure the validity of model fitting, the assumptions of linearity and normal, homoscedastic residual distributions were inspected. Effect size and estimated marginal means were employed in pairwise post-hoc testing and achieved with the emmeans package (Lenth, 2022). Cohen’s d effect size is employed and represents the difference in means as a function of the standard deviation. Commonly used interpretations separate effects sizes intro three categories; small (d = 0.2), medium (d = 0.5), and large (d = 0.8). Estimated marginal means represent the means predicted from the statistical model for each relevant independent variable combination. They allow comparisons between the dependent variables of interest, while accounting for the appropriate fixed and/or random effects in the model. Significance was set at α = 0.05 and pairwise and multiple comparisons were corrected using Tukey’s corrections for multiple comparisons.

## Results

In the efforts detailed here, we sought to understand and highlight instances where intrinsic properties of motor command integration introduce potential impediments in human force control. Persistent inward currents (PICs), an intrinsic property of spinal motoneurons, introduce a potent monoaminergic-dependent amplification and prolongation of excitatory synaptic currents that likely requires task-dependent inhibitory control. To highlight this relation, we designed two experimental paradigms that introduced potential constraints and conflicts to inhibitory control (see methods for rationale) and observed the subsequent alteration in estimates of PICs and control of force in human subjects. All tasks were performed about the ankle with PICs estimated from motor units (MUs) decomposed via high-density surface electromyography of the ankle dorsiflexors and plantar flexors.

### Experiment 1: Constraining inhibitory control degrades torque control

In the first paradigm, we forced a conflict between the inhibitory input necessary to deactivate PICs and continued excitatory drive to the motor pool. Figure 1a shows an example trial of this contraction paradigm, with the black trace indicating ankle torque and the discharge rate of decomposed motor units categorized and offset in the y-axis based upon their recruitment. For this contraction, individuals were asked to perform an initial 10% MVT stabilizing task (*plateau one)*, followed by a linear increase and decrease in torque (ramp), and a final 10% MVT stabilizing task (*plateau two)*. We refer to this task as a “sombrero”. As indicated in Figure 1a, MUs categorized as *brim* MUs were recruited at the onset of the stabilization task in plateau one, button MUs were recruited and derecruited in the center ramp region, and *cap* MUs were recruited in the center ramp but sustained discharge into the second plateau. Across participants, we decomposed 1327 MUs for the TA and 1157 MUs for the MG. The relationship between recruitment and derecruitment for the three categories of MUs can be appreciated in Figure 1b (left hand panel). Button MUs tend to be recruited and de-recruited at similar torque levels, whereas cap and brim units tend to be de-recruited towards the end of the contraction irrespective of recruitment thresholds.

**Figure 1:**
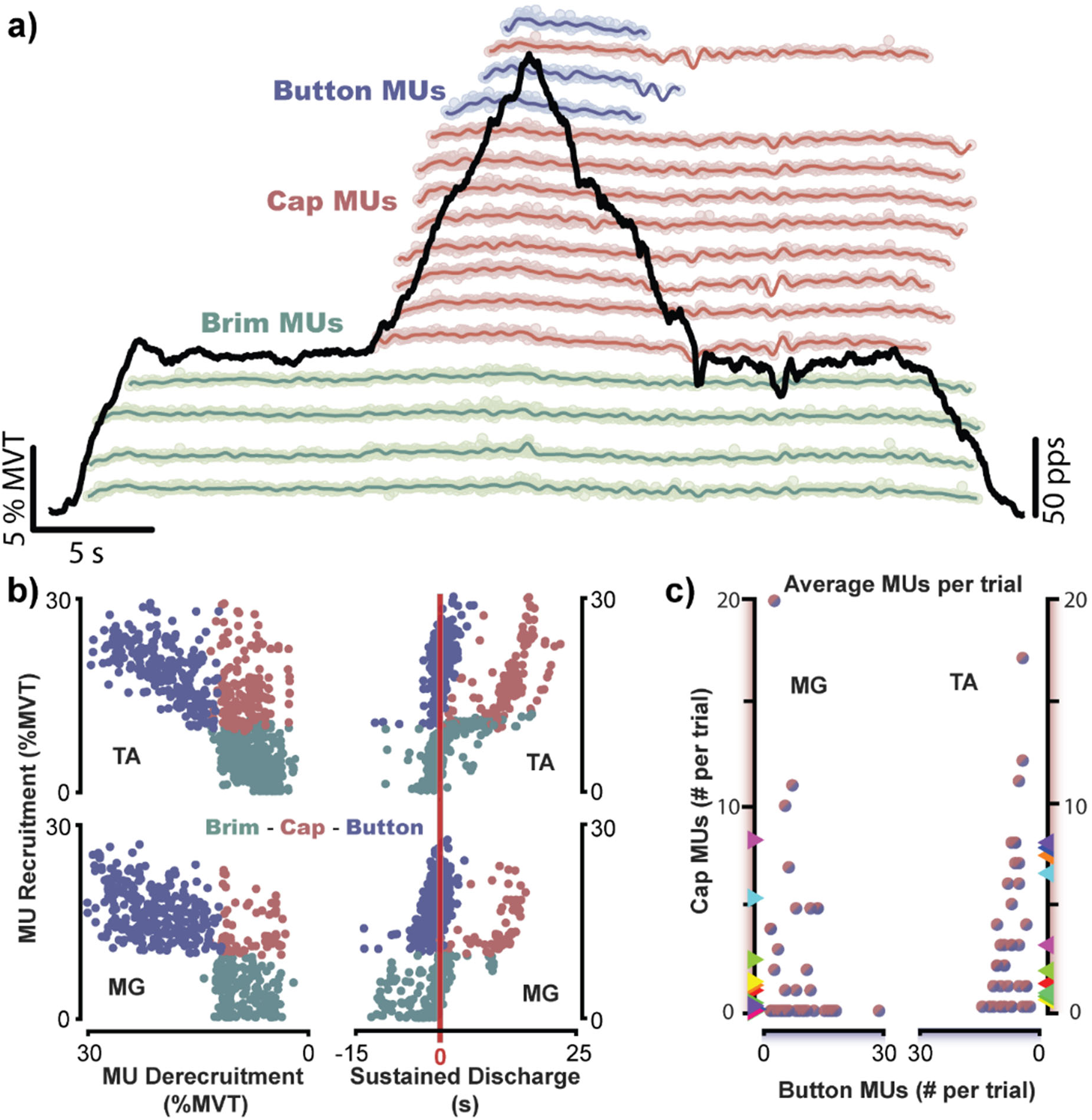
The sombrero contraction: a superimposition task that constrains inhibitory control of PICs. a) An exemplar sombrero contraction from a single participant, with dorsiflexion torque shown in the black trace normalized to an individual’s maximum voluntary torque (MVT). A collection of decomposed motor units (MUs) is shown smoothed and offset in the y-axis based upon when they were recruited in the contraction and colored according to their categorization. MUs categories included brim MUs that were recruited at the onset of the stabilization task in plateau one, button MUs that were recruited and derecruited in the center ramp region, and cap MUs which were recruited in the center ramp but sustained discharge into the second plateau. b) MU recruitment and derecruitment characteristics in the left column and duration of sustained discharge in the right column, with each dot representing an independent MU. Sustained discharge is represented as duration of time past theoretical derecruitment and the vertical red line indicates 0 s (i.e., a MU with identical recruitment and derecruitment torque). c) The average number of cap MUs as a function of button MUs decomposed per trial. Colored triangles along the y-axis represent participant averages.

During the sombrero contraction, a non-trivial number of MUs are recruited at levels greater than 10% MVT in the ramp contraction but not derecruited until the end of the second plateau (i.e., cap MUs). Across both muscles, an average of 10.35 MUs (95%CI: [8.00 12.63]) were recruited in the center ramp for a given trial, with trials that possess cap MUs displaying an average of 5 MUs (95%CI: [3.83 6.54]) sustaining discharge greater than two seconds into the second plateau (Figure 1c). The duration of sustained discharge past an expected theoretical derecruitment is shown in the rightmost column of Figure 1b (right hand panel). This duration is predicted by the category of unit (χ^2^(2) = 1038.2, p < 0.001) and is significantly greater in cap MUs than brim/button MUs (p < 0.001), with cap MU sustained discharge estimated as 10.91 s (95%CI: [9.46 12.37]) and 9.51 s (95%CI: [7.94 11.08]) for the TA and MG, respectively (Figure 1b).

The sustained discharge of cap MUs creates a situation where higher threshold MUs actively discharge during the second plateau. Interestingly, despite these cap MUs, torque output was close in magnitude between plateaus, as 10% MVT was demanded for both. Between plateaus we observed an average increase of 0.28% MVT (95%CI: [0.19 0.37]). Nonetheless, task performance was significantly degraded in the second plateau, as evidenced by the significant increase in coefficient of variation (CV) in ankle torque (Figure 2, left side). We found the plateau order to be predictive of CV (χ^2^(2) = 49.72, p < 0.001), with CV estimated to increase during *plateau two* by 1.73 % (95%CI: [1.21 2.24, d = 1.31]) for dorsiflexion trials and by 1.51 % (95%CI: [1.00 2.02], *d* = 1.15) for plantarflexion trials. These increases are notable when compared to the magnitude of CV during *plateau* one, increasing by more than 1.5-fold (plantar flexion: 2.60% [2.06 3.13]; dorsiflexion: 3.17% [2.62 3.71]). This increase in variation and apparent task difficulty can additionally be appreciated in the exemplar trials shown in Figure 1a and Figure 2. Despite demanding precise and accurate ankle torque, individuals were more variable in plateau *two*.

**Figure 2:**
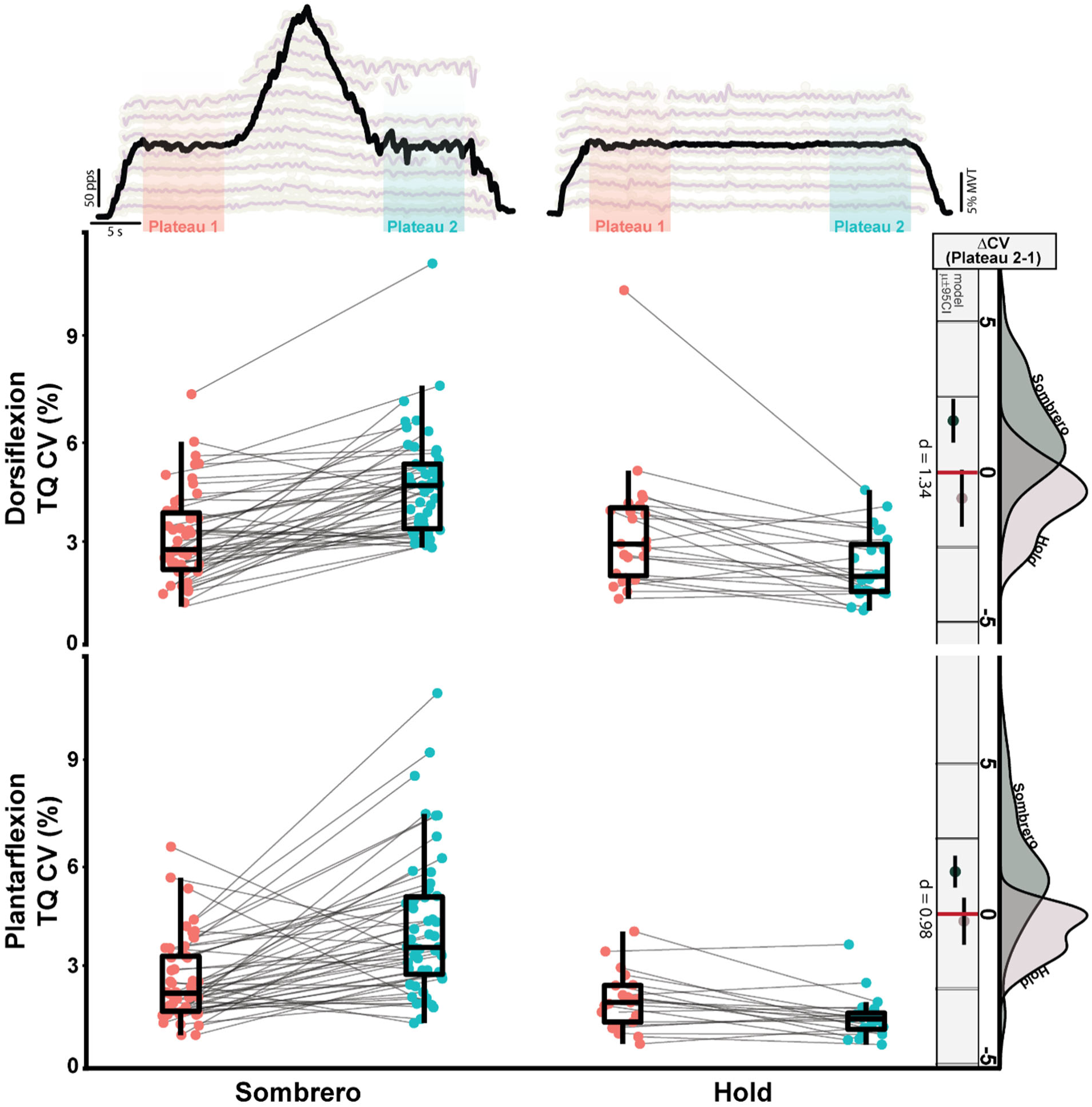
Torque variability increases during the second plateau in the sombrero contraction. Shown across the top are exemplar sombrero and hold contractions, with indicators for the first and second plateau region. (plateau 1 is red, plateau 2 is blue). The center panel represents the coefficient of variations (CV) in ankle torque for each trial in the first and second plateau region. Lines connect trials. The rightmost column represents the change in CV between plateaus (CV @ plateau 2 – CV @ plateau 1). The rightmost distributions indicate the probability density of these values for either the sombrero or hold contraction. The vertical line and colored circles indicate the model estimated mean value and 95% confidence interval for each contraction, with Cohen’s *d* effect size between the two contractions.

The increase in torque variability and greater number of higher threshold MUs in the second plateau is accompanied by a lower average MU discharge rate (Figure 3). We found plateau order to be predictive of average MU discharge rate (χ^2^(2) = 607.24, p < 0.001) with MUs actively discharging in the first plateau (i.e., brim MUs) lowering their average discharge rate in the second plateau by an estimated 1.75 pps (95%CI: [1.01 2.49]) and 2.40 pps (95%CI: [1.73 3.08]) for the TA and MG, respectively. The change in average discharge rate from *plateau one* to *two* is shown in Figure 3, with grey lines connecting unique MUs.

**Figure 3:**
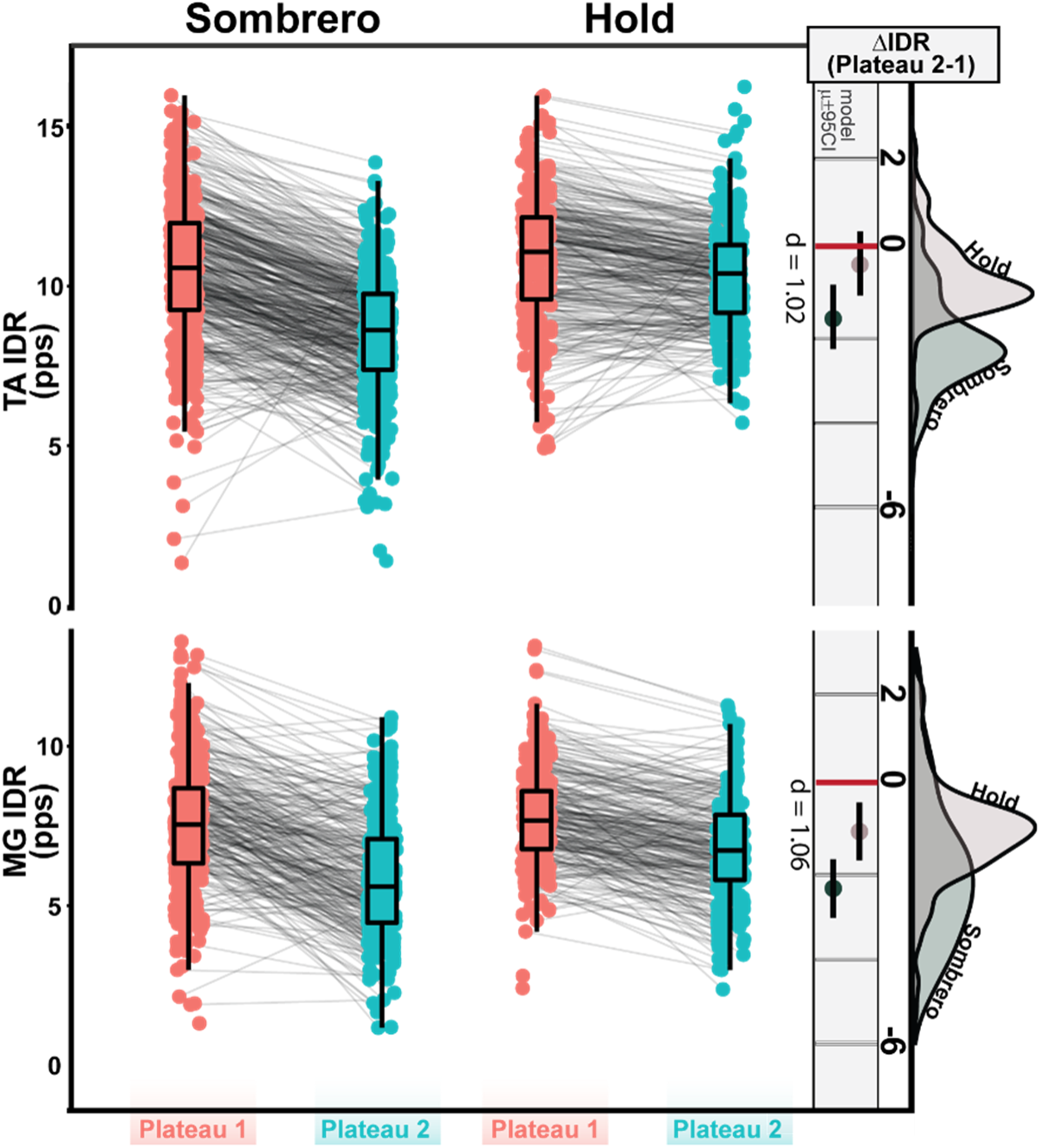
Average MU discharge rate decreases during the second plateau in the sombrero contraction. The center panel represents the average MU discharge rate for each trial in the first and second plateau region. Lines connect independent MUs. The rightmost column represents the change in discharge rate between plateaus (CV @ plateau 2 – CV @ plateau 1). The rightmost distributions indicate the probability density of these values for either the sombrero or hold contraction. The vertical line and colored circles indicate the model estimated mean value and 95% confidence interval for each contraction, with Cohen’s *d* effect size between the two contractions shown. (MG: Medial Gastrocnemius; TA: Tibialis Anterior; IDR: Instantaneous discharge rate)

To ensure that the observed MU characteristics and torque deficits were not the results of a time dependent habituation or a product of spike frequency adaptation, we additionally asked individuals to perform long stabilizing hold contractions (10% MVT). As expected, we found no significant increases in MUs in the second plateau and no regular occurrence of “cap” MUs. Furthermore, we found the order of plateau to significantly predict CV in ankle torque (χ^2^(2) = 18.35, p < 0.001) for the hold contractions, but found these values to decrease in *plateau two* for dorsiflexion (-1.04 % [95%CI: -1.81 -0.277], p = 0.008), and to not change for plantarflexion trials (p = 0.198). Also, we found no significant changes in MU discharge rate between *plateau one* and *two* for the TA (p = 0.153) and found a decrease of 1.08 pps (95%CI: [0.416 1.75], p = 0.004) in the MG (Figure 3).

Comparing the sombrero and hold contractions, we found the type of task to be predictive of both the change in torque CV (dorsiflexion: χ^2^(1) = 19.74, p <0.001 ; plantarflexion: χ^2^(1) = 11.96, p <0.001) and MU discharge rates (dorsiflexion: χ^2^(1) = 92.80, p <0.001 ; plantarflexion: χ^2^(1) = 73.33, p < 0.001), with the superimposition (i.e., sombrero) task eliciting greater increases in torque variation and decreases in discharge rates between the plateaus. Explicitly, brim MUs in the sombrero task show a significantly greater decrease in discharge rate from *plateau one* to *plateau two* when compared to stabilizing holds of the same duration and effort level (Figure 3, rightmost distributions) for both dorsiflexion (sombrero – hold: 1.24 pps [95%CI: 0.99 1.48]; *d* = 1.02) and plantarflexion (sombrero – hold: 1.34 pps [95%CI: 1.04 1.61]; *d* = 1.06). Furthermore, the second plateau of the sombrero contractions exhibit significantly greater increases in dorsiflexion (sombrero – hold: 2.59 [95%CI: 1.45 3.72], *d* = 1.34) and plantarflexion (sombrero – hold: 1.64 [95%CI: 0.69 2.59], *d* = 0.98) torque variability than the stabilizing holds (Figure 2, rightmost distributions).

### Experiment 2: Muscle length (joint angle) modulates PIC prolongation

To control the balance of excitatory and inhibitory input to motoneurons, we modulated the length of agonist-antagonist muscle pairs. Prior research has indicated ankle angle to acutely modulate the excitability of ankle muscle motor pools, such that spinal excitability is greatest as a muscle is shortened. Greater spinal excitability could indicate a relative disinhibition of motoneurons, thus reducing inhibitory constraints on PICs and potentiating PIC-induced prolongation of MU discharge. To observe this potentiation, we modulated the angle of the ankle (70°, 90°, and 110°) prior to linear isometric dorsiflexion and plantarflexion ramp contractions. The total number of decomposed MUs for each ankle configuration and muscle can be appreciated in Table 1.

**Table 1:** Maximum voluntary torque (MVT) and number of decomposed motor units (MUs) as a function of muscle length. Matched MUs indicate units that were decomposed at a minimum of two independent lengths. Values indicate mean and standard deviation.

|  | Long | Mid | Short |
| --- | --- | --- | --- |
| Maximum Torque (Nm) |  |  |  |
| Dorsiflexion | 24.38 (6.13) | 23.27 (6.31) | 15.62 (3.48) |
| Plantarflexion | 68.54 (26.63) | 57.38 (17.34) | 44.00 (11.24) |
| Decomposed MUs (#) |  |  |  |
| TA | 536 | 748 | 751 |
| MG | 641 | 698 | 637 |
| Matched MUs (#) | *Matched*<br>*to mid or short* | *Matched*<br>*to long or short* | *Matched*<br>*to mid or long* |
| TA | 153 | 222 | 93 |
| MG | 66 | 77 | 37 |

While we sought to modulate the balance of excitatory and inhibitory input to agonist motoneurons, changing the ankle angle, and subsequently agonist muscle length, also alters muscle force production capacity. To account for alterations in biomechanical properties (e.g., moment arm) and cross bridge dynamics of the agonist muscle we employed relative efforts, normalized to an individual’s maximal abilities for a given ankle configuration. This yielded lower maximal torque generation for shorter muscles (F = 24.09, p<0.001). That is, lower for dorsiflexion as the dorsiflexors were shortened (70°) and lower maximal plantarflexion torque as the plantar flexors were shortened (110°). These trends and quantitative values can be found in Table 1, with all isometric torque ramps normalized such that each was 30% MVT relative to the ankle configuration.

Figure 4b shows all decomposed MUs for the MG (plantarflexion trials) and TA (dorsiflexion trials) at each of the three ankle angles. For ease of interpretation, the angles have been categorized based upon the length of the agonist muscle during the contraction (i.e., 70° - long MG, short TA; 110° - short MG, long TA). For each respective plot, the smoothed discharge rate of individuals MUs are offset in the y-axis and colored according to the torque at which they were recruited with the black trace indicating the average torque performed across all trials and participants. A prolongation of MU discharge is readily apparent in this figure, with a greater number of teal higher threshold MUs that sustain discharge until the end of the ramp contraction when the muscle is in a shortened position. This indicates the expected shift in excitability and reduced ability to terminate PICs.

To quantify prolongation of MU discharge from PICs, we employed a paired MU analysis method (ΔF). Across both muscles, for MUs matched between at least two lengths, we found ΔF to be significantly predicted by muscle length (χ^2^(4) = 33.24, p < 0.001), with an estimated increase of 1.08 pps (95%CI: [0.47 1.69]) from long to short muscle lengths. Separating by muscle, from the longest to the shortest length, ΔF is estimated to increase by 1.29 pps (95%CI: [0.24 2.35]; *d* = 0.99) in the MG and 0.86 pps (95%CI: [0.24 1.48]; *d* = 0.66) in the TA (Figure 4c). To characterize prolongation from PICs and its contribution to torque generation, we quantified the duration of discharge that a MU exhibits on the descending phase of each ramp and the torque that each MU was derecruited at (Figure 4a). Across both muscles, we found the descending duration to be significantly predicted by muscle length (χ^2^(4) = 69.58, p < 0.001), with an estimated increase of 1.45 s (95%CI: [1.03 1.88]) from long to short muscle lengths. Separating by muscle, from the longest to the shortest length, descending duration is estimated to increase by 2.19 s (95%CI: [1.46 2.91]; *d* = 1.82) in the MG and 0.73 s (95%CI: [0.29 1.16]; *d* = 0.61) in the TA (Figure 4c). This indicates greater prolongation from PICs with MUs sustaining discharge for greater durations at shorter muscle lengths. Given that the MU discharge was prolonged at shorter lengths, the average derecruitment TQ of these MUs was decreased. We found the average torque at derecruitment to be significantly predicted by muscle length (χ^2^(4) = 64.94, p <0.001), with an estimated decrease from the longest to the shortest length by 5.04% MVT (95%CI: [3.28 6.79]; *d* = 1.74) for the MG and 1.80% MVT (95%CI: [0.75 2.86]; *d* = 0.62) for the TA (Figure 4c).

**Figure 4:**
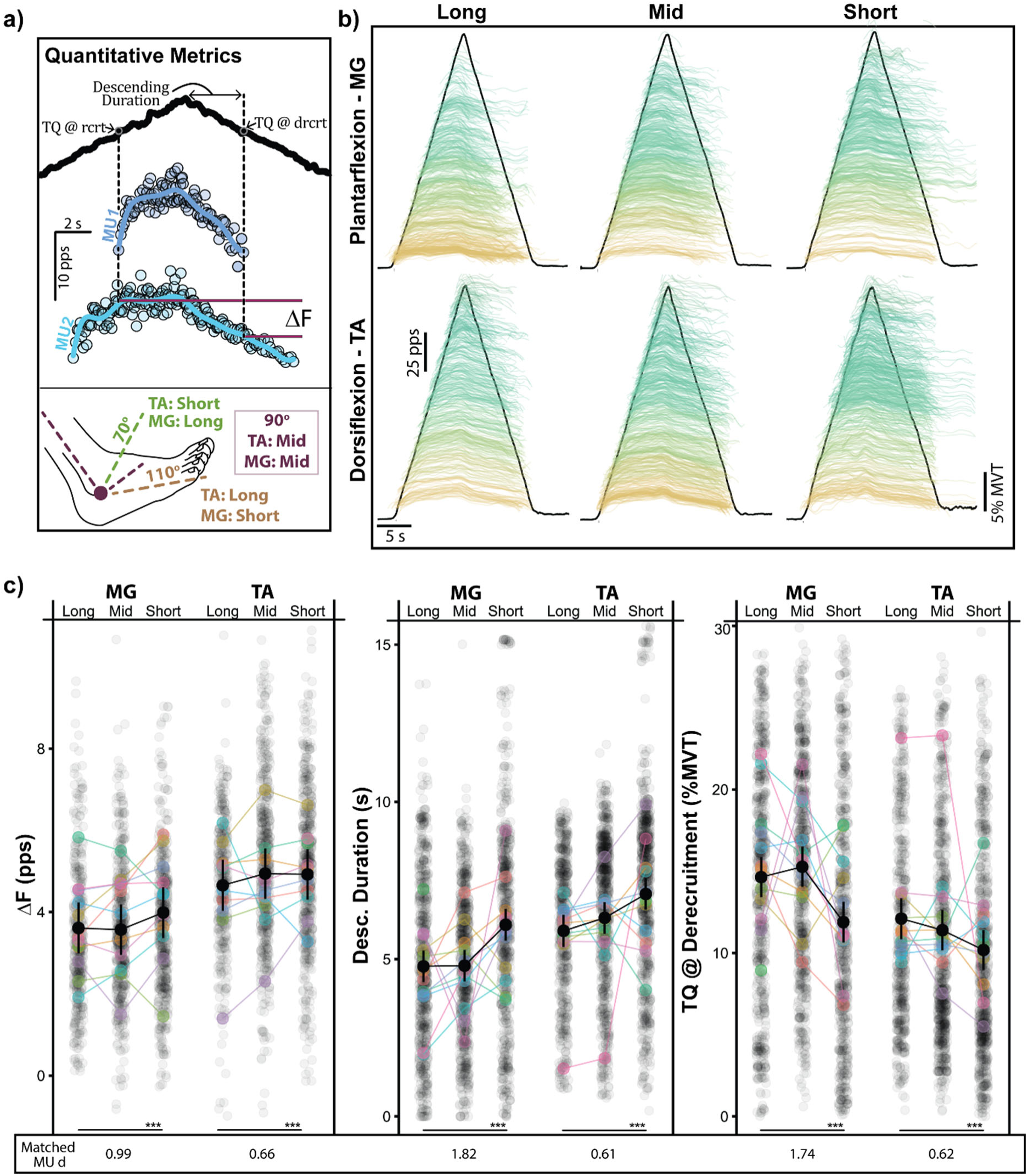
Prolongation of discharge from PICs is greatest at short agonist muscle lengths. a) The primary measures of PIC-induced prolongation are shown quantified for “MU1”. The employed ankle angles, and corresponding muscle lengths for the tibialis anterior (TA) and medial gastrocnemius (MG), are shown in the bottom of panel (a). In (b), all decomposed motor units (MUs) are shown for the TA and MG at each muscle length. MUs are smoothed and both colored and offset in the y-axis based upon when they were recruited in the triangular contraction. The average dorsiflexion or plantarflexion torque across all trials is indicated by the black trace. The quantitative metrics of prolongation from PICs are shown for the entire population of decomposed MUs at each of the three lengths in (c). This is the paired MU analysis (ΔF), the duration of time spent on the descending (desc.) portion of the ramp, and the torque (TQ) at which a MU is derecruited. Additionally, the effect size (Cohen’s *d*) between the differences observed at the long and short lengths is shown across the bottom for matched MUs decomposed at a minimum of two independent lengths.

Of note, although the data depicted in Figure 4b and Figure 4c include the entire dataset, the primary statistics that we report are based on only those MUs matched at a minimum of two independent lengths (see Methods: *MU Matching*). The trends displayed in Figure 4 remain with the matched MUs and all metrics for the matched MUs can be observed in the supplementary information.

### Experiment 3: Greater PIC prolongation at short muscle lengths further degrades torque control

Given the degradation of torque control observed alongside greater PIC prolongation in experiment one and the modulation of PIC prolongation with muscle length in experiment two, we sought to show that changes in muscle length could manipulate the observed degradation in torque control. If PIC prolongation does indeed generate the amplified torque variability observed in experiment one, facilitating this behavior through shortening an agonist muscle should further degrade torque control, and vice versa. To investigate this relation, we had individuals perform both the superimposition task as well as the long stabilizing hold contractions of experiment one (see Figure 2) at the two extreme ankle joint angles (70° and 110°) tested in experiment two. When short, we decomposed 1123 MUs for MG and 1051 MUs for the TA while when long we decomposed 1005 MUs for MG and 908 MUs for the TA.

A stark increase in PIC prolongation behavior can be appreciated in Figure 5a at shorter muscle lengths, with brim, cap, and button MUs colored as indicated. A clear increase in the number of cap MUs is observed at shorter lengths, with their continuous discharge estimates shown in red and often lasting through the second plateau. Across both muscles, length was found to significantly predict the number of cap MUs (χ^2^(4) = 55.54, p <0.001), with an estimated 5.17 (95%CI: [2.60 7.75]) more cap MUs per trial for the MG and 4.41 (95%CI: [1.75 7.10]) per trial for the TA (Figure 5d) at shorter lengths. Additionally, the duration of sustained discharge for these cap MUs can be observed in Figure 5b, with the proportion of units that sustain discharge for more than 2 s into the second plateau significantly greater at shorter lengths (χ^2^(2) = 49.09, p <0.001) for both dorsiflexion (*d* = 0.98) and plantarflexion (*d* = 1.38).

**Figure 5:**
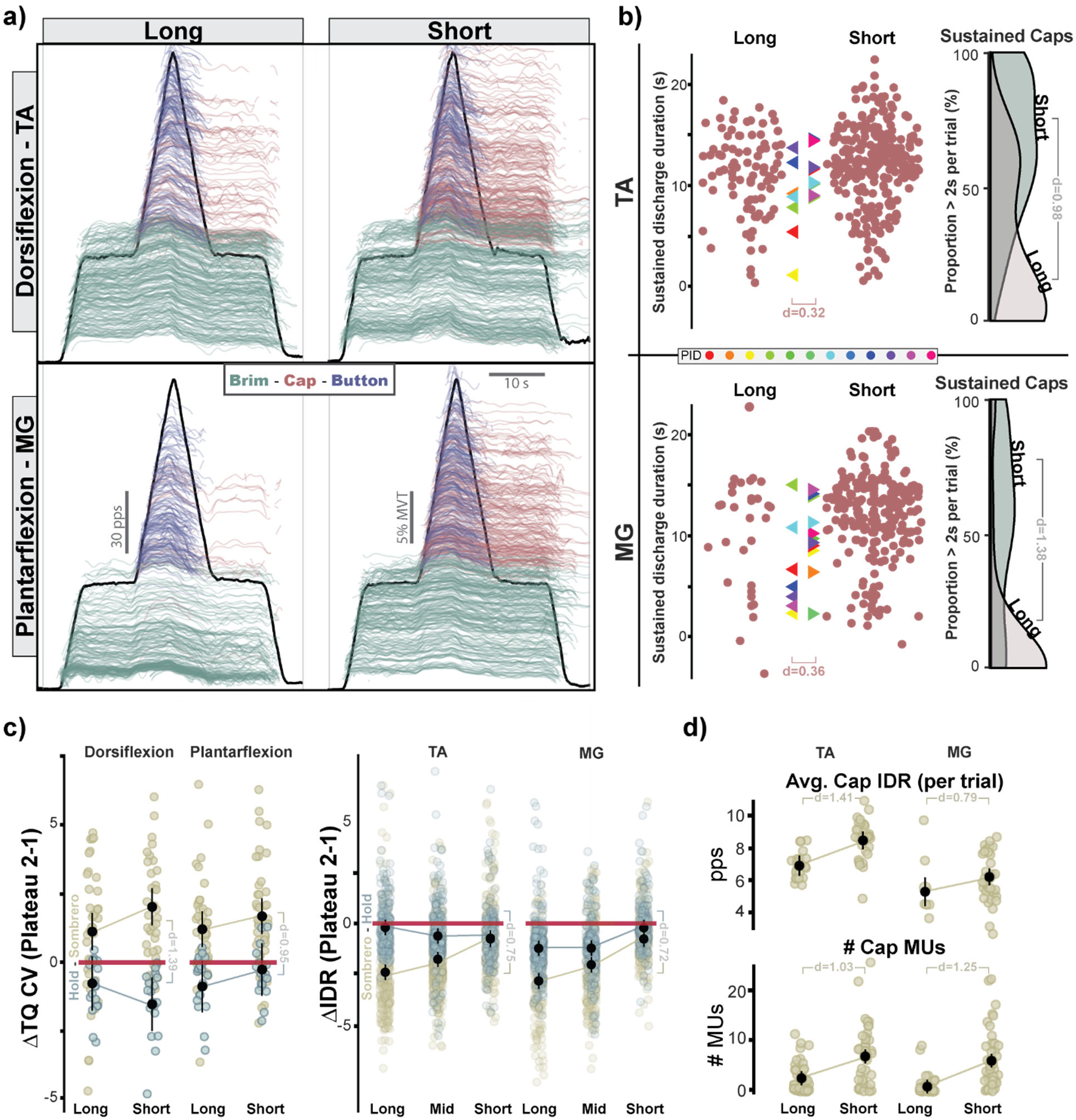
Greater PICs and torque variability at shorter muscle lengths during the sombrero contraction. Shown in (a) is the entire population of decomposed motor units (MUs) for the tibialis anterior (TA) and medial gastrocnemius (MG) at long and short muscle lengths. MUs are smoothed and offset in the y-axis based upon when they were recruited in the contraction. The average dorsiflexion or plantarflexion torque across all trials is indicated by the black trace. MU categories are indicated by colors. In the left portion of (b), the duration of sustained discharge past a theoretical derecruitment is shown for all cap MUs, with colored triangles indicating participant averages. In the rightmost portion of (b), probability density distributions for the proportion of MUs that turn on in the center ramp that sustain discharge 2 seconds (s) into the second plateau are shown for both lengths. The change in CV (plateau 2 – plateau 1) is shown in the left panel of (c) for the sombrero and hold contractions at the long and short lengths. In the right panel of (c), the average change in brim MU discharge rate (plateau 2 – plateau 1) is shown. The black vertical lines and dots represent the model estimated means and 95% confidence interval for each contraction and length. In (d) the average cap MU discharge rate for each trial is shown in the top row and the number of cap MUs for each trial is shown in the bottom row.

In tandem with greater cap MUs at shorter muscle lengths, we additionally found task performance in the second plateau to further degrade at shortened muscle lengths. When comparing the increase in CV in ankle torque from *plateau one* to *plateau two* (Figure 5c; ΔTQ CV), we found muscle length to be a significant predictor of this increase in variability (χ^2^(1) = 6.169, p = 0.013, *d* = 0.39) across muscle. Separating by muscle, we found shorter muscle lengths to produce the highest increases in CV, with estimated increases of 2.00 % (95%CI: [1.24 2.76]) for dorsiflexion and 1.70 % (95%CI: [0.98 2.42]) for plantarflexion. In comparison, at the longer muscle lengths CV in ankle torque was increased by an estimated 1.13 % (95%CI: [0.37 1.89]) for dorsiflexion and 0.84 % (95%CI: [0.10 1.58]) for plantarflexion.

Interestingly, when investigating the average discharge rate of MUs during the second plateau, though the general trend of lower average discharge rates in plateau two remained, this change was greatest at long muscle lengths. In specific, despite a greater number of cap MUs actively discharging in plateau two at shorter muscle lengths, we found brim MU discharge rate to decrease by less between plateaus and found greater cap MU discharge rates. This can be observed in Figure 5c for the change in brim motor unit discharge rate between plateaus and in Figure 5d for the average cap MU discharge rate. Across muscles, we found the change in brim MU discharge rate to be significantly predicted by muscle length (χ^2^(4) = 521.12, p < 0.001), with shortest lengths producing the lowest decrease between plateaus (Long - [MG: 2.78 pps [95%CI: 2.28 3.27], TA: 2.40 pps [95%CI: 1.92 2.88]; short - [MG: 0.803 pps [95%CI: -0.32 1.29], TA: 0.70 pps [95%CI: 0.22 1.18]). Furthermore, we found cap MU discharge rate to be significantly predicted by muscle length (χ^2^(2) = 23.16, p < 0.001), with average discharge rate estimated to increase by 1.26 pps (95%CI: [0.69 1.84]) across the MG and TA at shorter compared to longer muscle lengths.

To estimate common drive to the motor pool and its potential relation to cap MUs and degradation in torque control, we conducted a coherence analysis of MUs actively discharging in either plateau. The change in average low frequency coherence (0-5 Hz) can be seen in Figure 6, with each data point representing an individual trial. Comparing the change between plateaus, we found muscle length to be a significant predictor (χ^2^(4) = 39.21, p < 0.001). For long muscle lengths, average coherence is estimated to change by -0.63 (95%CI: -1.00 -0.25]) and 0.08 (95%CI: -0.30 0.46]) for the MG and TA, respectively. In contrast, at short lengths, average coherence is estimated to increase by 0.67 (95%CI: [0.28 1.07]) in the MG and 0.69 (95%CI: [0.31 1.08]) in the TA. To determine the potential relation of this shared common drive to ankle torque fluctuations, we ran a similar analysis between actively discharging MUs and the resultant ankle torque. Looking at the change in average low frequency coherence between MUs and torque, we saw a similar trend with muscle length a significant predictor of this coherence (χ^2^(4) = 29.09, p < 0.001).

**Figure 6:**
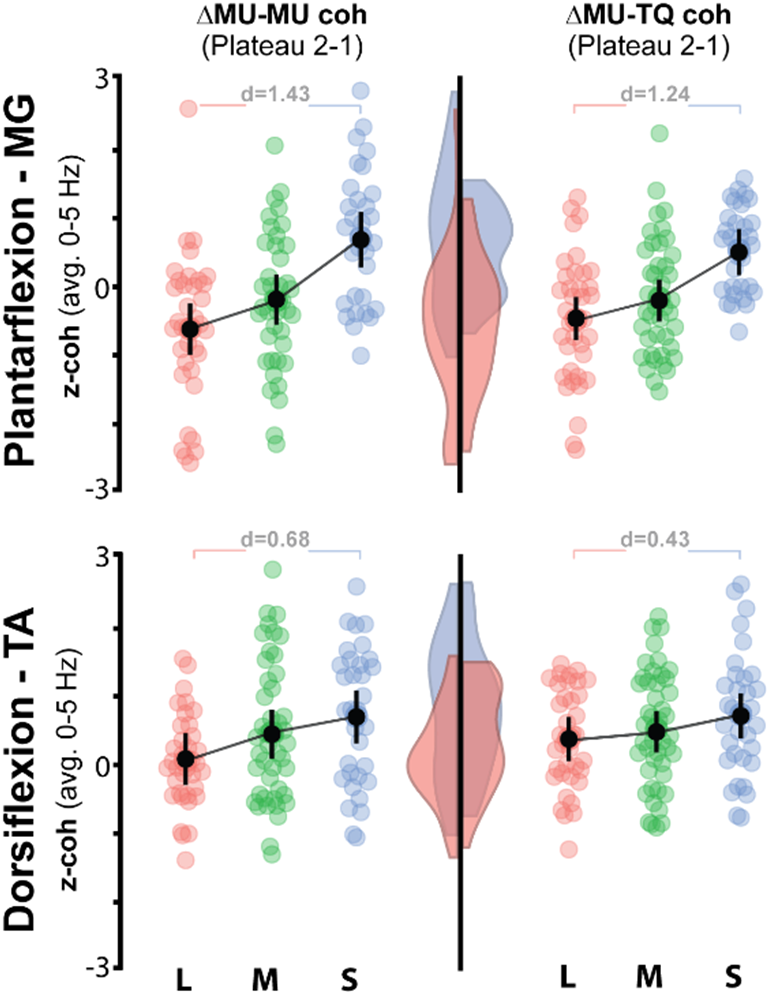
Coherence between motor units (MUs) and between MUs and torque (TQ) is increased at short muscle lengths. Shown is the change in average low frequency z-transformed coherence (0-5 Hz) between plateaus of the sombrero contraction (plateau 2 – plateau 1). This is shown for coherence between MUs on the left and between MUs and ankle TQ on the right. The probability density distributions for these values are shown for the long (pink) and short (blue) overlayed and projected on the center vertical line. Within plots, each dot represents a trial and black vertical lines and dots represent the model estimated means and 95% confidence interval for each contraction and length.

As with experiment one, to ensure that these observed MU characteristics and torque deficits were not the results of a muscle dependent habituation or phenomena, we additionally asked individuals to perform long stabilizing hold contractions at each ankle angle. Looking at these sustained hold contractions, we unsurprisingly found no significant increases in MUs in the second plateau or regular occurrence of cap MUs dependent on muscle length.

Comparing the stabilizing hold and sombrero contractions, we found sombreros to exhibit significantly greater variation in ankle torque (χ^2^(4) = 66.98, p < 0.001) for both dorsiflexion (sombrero-hold: Long 1.93% [95%CI: 0.46 3.40]; Short 3.59% [95%CI: 2.12 5.06], *d* = 1.39) and plantarflexion (sombrero-hold: Long 1.79% [95%CI: 0.38 3.19]; Short 1.97% [95%CI: 0.55 3.39], *d* = 0.95), but no interaction between the contraction type and length (χ^2^(3) = 7.12, p = 0.07). Furthermore, we found the change in brim MU discharge rate between plateaus to be predicted by contraction type (χ^2^(6) = 431.22, p < 0.001), being lower during the sombrero for all but the shortest muscle length for dorsiflexion (sombrero-hold: Long -2.21 [95%CI: -2.49 -1.94]; Mid -1.11 [95%CI: -1.35 -0.87]; Short -0.06 [95%CI: -0.23 0.35], *d* = 0.75) and plantarflexion (sombrero-hold: Long -1.60 [95%CI: -1.87 -1.32]; Mid -0.86 [95%CI: -1.13 -0.59]; Short -0.69 [95%CI: -0.96 -0.41], *d* = 0.72).

## Discussion

In this study, we showcase instances where intrinsic properties of motoneurons introduce potential impediments to human motor control. Dendritic PICs in motoneurons introduce a monoaminergic dependent amplification and prolongation of excitatory synaptic inputs that requires task dependent control, less they engender aberrant discharge patterns and poorly controlled forces. In support of this, we show through a series of paradigms that challenging the inhibitory control of PICs exacerbates prolongation of motoneuron discharge and introduces subsequent problems in torque control.

### Prolongation of discharge from PICs degrades torque control

In the first paradigm of this study, we asked individuals to perform a superimposition task (i.e., sombrero contraction) that challenged the inhibitory input necessary to deactivate PICs (for rationale, see Methods: *Experiment 1*). In short, this superimposition task was comprised of an isometric ramp atop a low-effort stabilizing hold, where individuals were to perform the ramp contraction and return to the stabilizing hold as precisely as possible (see Figure 1a). Upon returning to the hold (plateau 2), the inhibitory input necessary to deactivate the PICs of the MUs recruited for the ramp contraction was compromised as continued excitatory drive is necessary to accomplish the stabilizing hold. Subsequently, many of these newly recruited MUs sustained discharge (i.e., prolongation) and became what is termed cap MUs (Figure 1a). This created a situation where higher threshold MUs were actively discharging in the second plateau, which forced the entire motor pool to discharge at a lower average rate (Figure 3) and introduced difficulties in control (Figure 2).

The notion that the cap MUs, and thus prolongation from PICs, is responsible for the degradation in torque control in plateau two is supported by our findings that stabilizing holds of the same duration did not exhibit this phenomenon and the findings of the third experiment where modulating PIC prolongation also modulated torque variability. Regarding the former, when comparing the superimposition task and stabilizing holds, the superimposition task exhibited a significantly greater reduction in average MU discharge rate and a significant increase in the coefficient of variation (CV) in torque. Comparing the two contractions, the change in CV from plateau one to plateau two is estimated to be greater in the sombrero contraction by 2.6% in dorsiflexion and 1.6% in plantarflexion. Given the average mid length plateau one CV was found to be 2.60% for plantarflexion and 3.17% for dorsiflexion, this increase is quite substantial and likely represents a meaningful degradation in control (∼100% increase). Of note, force steadiness and CV is thought a proxy for neural drive to the motor pool and often considered predictive of motor function in health and disease (Enoka and Farina, 2021, Oomen and van Dieen, 2017).

### Muscle length (joint angle) modulates prolongation from PICs

Altering ankle joint angle, and subsequently muscle length, has long been appreciated to modulate measures of spinal reflex excitability, with shorter ankle muscles producing greater electrophysiological estimates of spinal excitability (Burke et al., 1983, Gerilovsky et al., 1989, Gerilovsky et al., 1977). In specific, in addition to greater Mmax and Hmax at shorter muscle lengths, both plantar and dorsiflexor muscles exhibit greater Hmax/Mmax ratios, indicating greater spinal reflex pathway excitability that peripheral mechanisms of the muscle alone cannot account for (Dutt-Mazumder et al., 2020, Frigon et al., 2007, Hwang, 2002, Patikas et al., 2004). Explanations and discussions regarding these observations are varied and have included altered motoneuron excitability, presynaptic inhibition of Ia afferents, muscle fiber volume, golgi tendon organ (Ib) feedback, and the relation of muscle fibers to the skin (Garland et al., 1994, Gerilovsky et al., 1989, Frigon et al., 2007, Hwang, 2002, Patikas et al., 2004).

We speculate that the observed changes in prolongation in the second experiment with changes in joint angle are due to changes in inhibitory input to motoneurons. This speculation is based upon our observations that MUs have longer average durations of discharge on the descending limb of the ramp and lower average derecruitment torques at shorter muscle lengths for both the entire decomposed MU population (Figure 4c) and for MUs identified during at least two length conditions (i.e., matched MUs; supplementary Figure 1). Analyzing the discharge trajectories of matched MUs indicates a rightward shift in MU activity, with later recruitment and derecruitment and increased time on the descending phase.

PICs are particularly sensitive to inhibitory inputs, in part due to their dendritic location, being deactivated by both recurrent and reciprocal inhibition (Bui et al., 2008a, Hyngstrom et al., 2007, Hyngstrom et al., 2008, Bui et al., 2008b, Kuo et al., 2003). Importantly, the control of PICs is likely only achieved via inhibitory inputs, as excitatory inputs (e.g., Ia muscle spindle afferents) only activates PICs, which remain active even after cessation of this excitatory input (Heckman and Enoka, 2012). It is possible that changes in ankle angle could alter inhibitory input via multiple means to produce greater prolongation from PICs at short muscle lengths. In addition to altering agonist muscle length and discharge of spindle afferents, changes in ankle angle also alters antagonist muscle length (i.e., reciprocal inhibition), passive tension on the tendon, joint and cutaneous feedback, and heteronymous input from synergist muscles, all of which could alter synaptic input to agonist motoneurons and contribute to the observed results (Crone et al., 1987, Nichols, 2018, Hunt, 1952, Fallon et al., 2005, Pierrot-Deseilligny and Burke, 2012, Baxendale and Ferrell, 1982, Hongo et al., 1984).

Furthermore, since we employed relative efforts and the absolute forces required for the ramp contractions at each muscle length were necessarily different, golgi (Ib) feedback is likely altered and may explain the observed results. Though Ib feedback is certainly complex and less understood that Ia feedback (Pierrot-Deseilligny and Burke, 2012), given that the ramp contractions at shorter muscles lengths were performed at absolute values of torque lower in magnitude, inhibition from golgi tendon organs could be lower during these contractions (Houk et al., 1971, Jami, 1992). This would lower the overall inhibitory drive to motoneurons and allow for greater prolongation from PICs. Similarly, greater Ib inhibition at longer muscle lengths when both the absolute force generation for the contraction and the passive tendon/muscle tension is greater, could reduce the effective drive of the dendritic current of PICs.

Lastly, lowering the voltage threshold for PICs could explain the observed results. A lower PIC activation threshold would cause PICs to activate earlier and deactivate later in the ramp contractions for the same effort. Though MUs matched between conditions followed this general trend, they also displayed greater normalized torque at recruitment which would appear at odds with a lower PIC voltage threshold. That said, of critical importance, these represent relative torque values (i.e., normalized to max for each angle, see Table 1) which makes the absolute magnitude of torque values lower at shorter lengths. Thus, in an absolute sense, this makes the lower derecruitment torques more impressive and would diminish any reported increases in torque at recruitment.

### Further degradations in torque control with greater prolongation from PICs at short muscle lengths

In the third and final experiment of this study, we combined the superimposition task of the first experiment and the extreme changes in muscle length of the second experiment. Combining these paradigms, we again observed the prolonging behavior of PICs to be greatest at short muscle lengths (Figure 5a,b,d) and found this behavior to further exacerbate impairments to torque control (Figure 5c).

Through putting a muscle in a shortened positions, we found the propensity for sustained discharge of MUs recruited in the ramp portion of the sombrero to be markedly increased (see Figure 5a,b). When observing the proportion of these newly recruited MUs that sustained discharge more than 2s past their theoretically expected discharge instance, a large increase in the average proportion of MUs is observed comparing long to short lengths (*d* = 0.98, 1.38 for TA, MG). This simultaneously strengthens the observations of greater prolonging behavior at shorter lengths found in the second experiment while amplifying the negative ramifications of this behavior in the superimposition task from the first experiment.

We reason that the greater number of cap MUs actively discharging in the second plateau at short muscle lengths is responsible for this degradation of torque control observed in the sombrero task. To perform the center ramp portion of the sombrero contraction, higher threshold MUs are recruited which then subsequently sustain discharge into the second plateau (i.e., cap MUs). As descending excitatory drive to the motor pool is progressively decreased to perform the descending portion of the ramp, the sustained discharge in these cap MUs is likely maintained primarily by the intrinsic excitatory current from PICs. Since PICs are keeping cap MUs above their activation threshold, descending drive would similarly entrain these MUs and thus their discharge rates could represent oscillations in common drive to the motor pool. Interestingly, estimates of common drive to MUs were found to increase at shorter lengths, alongside a greater shared oscillation between the ankle torque being performed and the discharge of the motor pool (Figure 6). We speculate that this oscillation may introduce difficulties in control as variations in descending drive now entrain many more MUs, particularly a greater proportion of higher threshold MUs with relatively larger muscle twitches and greater force production capacity (Henneman and Mendell, 2011). In specific, oscillations in the discharge rate of these higher threshold cap MUs generate greater deviations in torque and this likely leads to complexities in control (e.g., over-corrections).

As was speculated and observed in the first experiment, the average discharge rate of the entire motor pool decreased during the second plateau of the sombrero contraction, likely to accommodate the greater number of actively discharging MUs. Interestingly, though brim units still generally exhibited a decrease in discharge rate from plateau one to two (negative values, Figure 5c), these values became progressively less negative at shorter muscle lengths. Similarly, the average discharge rate of cap MUs also displayed a relative increase as the muscle was put into shorter lengths. This is despite a greater number of MUs actively discharging at shorter lengths, with significantly greater cap MUs (Figure 5d) now recruited and present. Though seemingly peculiar, as now many more MUs are actively discharging at greater rates to produce the same relative (lower absolute) force, this is likely due to a complex interaction with the change in twitch properties of MUs. It has been shown that the twitch course of MUs possess a shorter duration and half-relaxation time constant when a muscle is shortened. Thus, at shorter lengths, the summation of twitches is likely less and higher discharge rates may be required to produce the same force (Bigland-Ritchie et al., 1992, Christova et al., 1998, Vander Linden et al., 1991, Rack and Westbury, 1969, Heckman et al., 1992).

### Implications: Neurological Impairment and Muscle Cramps

The findings presented here suggest that difficulties in human torque control arise when perturbations and constraints are placed on the inhibitory control of PICs. Though long appreciated for their critical role in motor control, PICs may introduce problems in motor control at the level of the motoneuron when improperly managed. This is particularly relevant to neurological conditions where pathological shifts in monoamines have been theorized (e.g., spinal cord injury, chronic stroke) (Li et al., 2019, Murray et al., 2010, McPherson et al., 2018b, Beauchamp et al., 2022a). Monoamines (i.e., norepinephrine and serotonin) amplify PICs by upwards of five-fold, markedly potentiating the amplification and prolongation of discharge by PICs. Thus, when monoaminergic drive to the cord is pathologically high, such as is theorized in chronic hemiparetic stroke {Beauchamp, 2022 #709}{McPherson, 2008 #63;McPherson, 2018 #542;McPherson, 2018 #87}, PICs will be magnified and make motoneurons hyperexcitable. These hyperexcitable motoneurons are much more sensitive to afferent inputs which may lead to spasticity and even hypertonicity, where individuals are unable to derecruit and cease discharge of motoneurons. Furthermore, more excitable motoneurons also amplify the commands of weak and indirect motor pathways (e.g., reticulospinal) which is theorized to generate a loss of independent joint control (Dewald et al., 1995, Ellis et al., 2016, Li et al., 2019, McPherson et al., 2018a, McPherson et al., 2018c, McPherson and Dewald, 2022).

In addition to neurological conditions, findings from the latter two experiments may provide an initial link between PICs and muscle cramps. As the reader may appreciate, muscle cramps, particularly in the calf (i.e., triceps surae) can be acutely debilitating. Surprisingly, generative and facilitating factors of muscle cramps are still debated. Though dehydration and ion concentration imbalances are often suggested, causative support for this is lacking and largely comprised of observational reports and medical records (Maughan and Shirreffs, 2019). That said, it has often been recognized that muscle cramps are more prevalent in shortened muscles and can be alleviated by transient forms of inhibitory inputs (Khan and Burne, 2007, Mills et al., 1982). Given the sensitivity of PICs to inhibitory inputs, and their ability to generate prolonged and sustained discharge in MUs, logic may suggest that PICs play a role in muscle cramps. Indeed, it has been speculated that PICs could contribute to this phenomenon, though a direct link has remained unestablished (Baldissera et al., 1991, Minetto et al., 2013, Khan and Burne, 2007). This study provides an initial link, demonstrating that the prolongation caused by PICs is most pronounced at shorter muscle lengths (Experiment 2) and that this effect may hinder voluntary derecruitment of MUs (Experiment 3), potentially resulting in sustained muscle contractions akin to those observed in muscle cramps. Additional studies in this realm are certainly warranted.

### Considerations and Limitations

Several considerations and limitations must be appreciated when interpreting this work. First, findings are necessarily limited by the population of MUs decomposed with HDsEMG and may not represent the entire motor pool. The decomposition of surface EMG is biased towards larger MUs and MUs closer the electrode (Farina and Holobar, 2016, Caillet et al., 2023). Additionally, as noted, changes in joint angle result in many changes in addition to changes in muscle length (i.e., afferent input, descending drive). Furthermore, though antagonist EMG was monitored throughout the contractions and no substantial deviations were observed, contribution from the antagonist to the observed results cannot be ruled out. Notably, this work fails to replicate similar work in the decerebrate cat. Studies from our group in animal preparations show that PICs in agonist motoneurons are markedly potentiated as a result of reduced reciprocal inhibition when its antagonist muscle is shortened (Hyngstrom et al., 2007). This is in direct contrast to the findings of this study and implies that motoneuron outputs would suffer from excessive PIC prolongation at long agonist muscle lengths. That said, these experiments were in the decerebrate cat which lacks descending corticospinal input and were not the result of volitional contraction as was employed in the present study. Of note, reciprocal Ia inhibitory post synaptic potentials in the baboon were found to only be evoked after spinal transection or depressed brain function, implying a descending tonic inhibition of reciprocal Ia afferents (Hongo et al., 1984). Perhaps humans exhibit similar descending inhibitory control, which could explain the divergence of our results from the decerebrate cat and potentially implicate changes in descending presynaptic Ia inhibition as a contributing mechanism. Indeed, a reduction in presynaptic Ia inhibition at shortened muscle lengths may be advantageous given the deprecated force production capacity of shortened muscles.

## Conclusion

This work explored how intrinsic electrical properties of spinal motoneurons, which transduce motor commands into muscle contraction, may introduce potential errors in control. In specific, motoneurons possess monoaminergic dependent persistent inward currents (PICs) which amplify and prolong excitatory synaptic inputs to motoneurons. These PICs play a pivotal role in controlling motor output, adapting motoneuron excitability to meet the diverse demands of motor tasks. However, not all tasks require uniform amplification and prolongation of excitatory inputs to motoneurons, and a precise inhibitory control mechanism is likely necessary. Thus, we devised two isometric contraction paradigms that challenged the inhibitory control of PIC behaviors and reasoned that these challenges would degrade ankle torque control. The first paradigm combined a discrete task with a low-effort stabilization task, compromising inhibitory input and resulting in excessive PIC prolongation and more variable torque control. The second paradigm altered spinal excitability by adjusting muscle length, hindering PIC deactivation, and leading to increased PIC prolongation at shorter agonist muscle lengths. Combining these paradigms further highlighted the difficulties that arise when control over PICs is disrupted, with the deficits in torque control observed in the first paradigm exacerbated at shorter muscle lengths when PIC prolongation is most pronounced. These findings underscore the pivotal role of task-dependent inhibitory control in managing PICs, with potential implications for clinical conditions characterized by pathological shifts in monoaminergic drive and the potential to shed light on mechanisms underlying muscle cramps.

---

**Figure S1:**
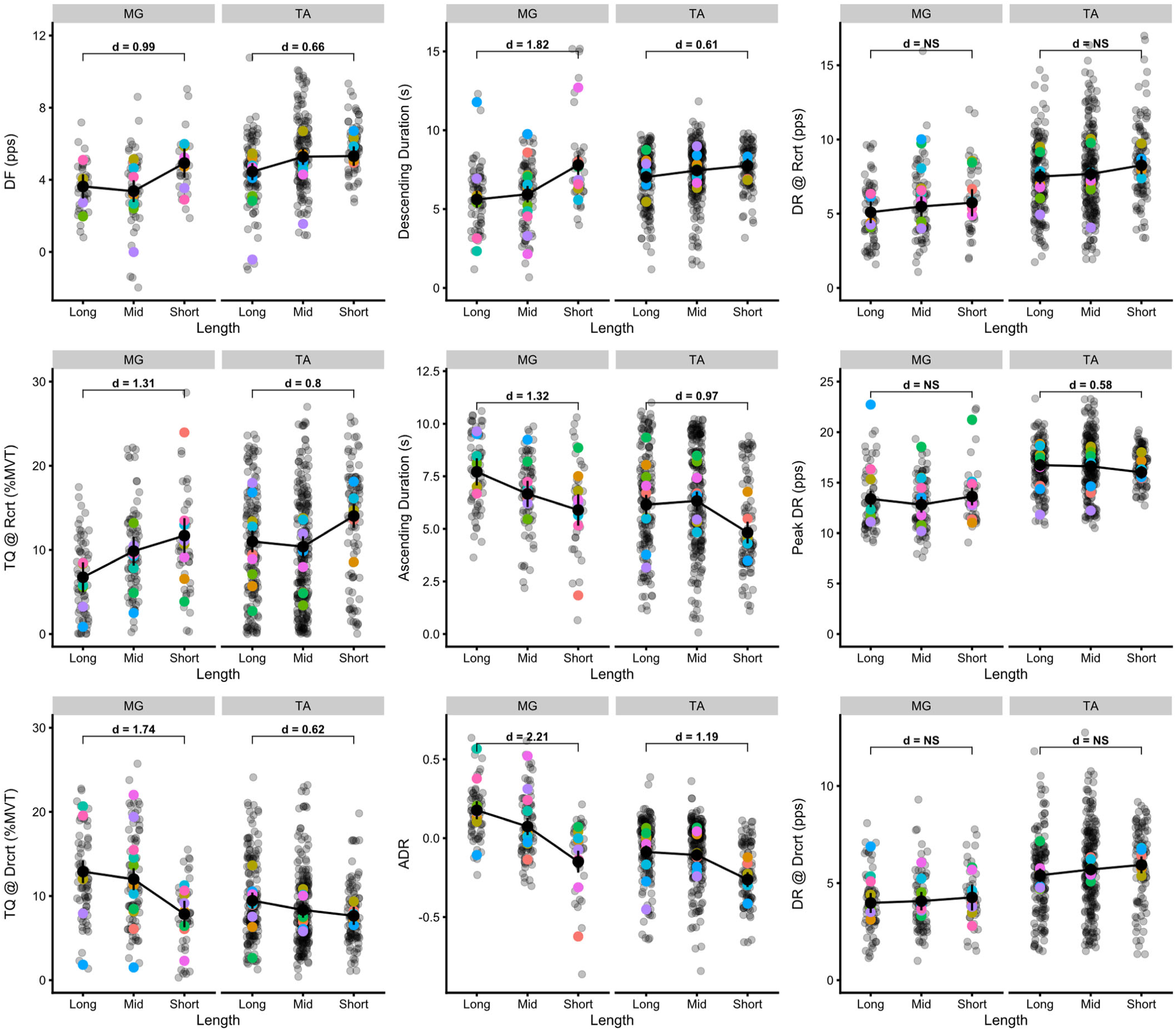
Discharge characteristics for motor units matched between lengths. The top row from left to right represent ΔF, the duration of time spent on the descending portion of the ramp, and discharge rate (DR) at motor unit recruitment (Rcrt). The middle row represents the torque (TQ) at motor unit recruitment, the time spent on the ascending portion of the ramp, and the peak discharge rate. The bottom row represents torque at derecruitment (Drcrt), the difference between ascending and descending duration as a function of the entire duration of discharge (ADR), and the discharge rate at derecruitment. Colored dots represent participant averages, grey dots represent raw data points for individual motor units, and the black connecting lines indicate the estimated marginal means predicted by the linear mixed effects model. Vertical black lines indicate the 95% confidence interval for these estimated marginal means. Cohen’s d effect size is shown for the difference between long and short when the differences are predicted as significant by the mixed model (p<0.05) and are quantified with the estimated marginal means. Significance for the fixed factor of length and its interaction with muscle is as follows: ΔF: (χ^2^(4) = 33.24, p < 0.001); Descending Duration: (χ^2^(4) = 69.58, p < 0.001); DR @ Rcrt: (χ^2^(4) = 7.74, p = 0.101); TQ @ Rcrt: (χ^2^(4) = 77.07, p < 0.001); Ascending Duration: (χ^2^(4) = 90.33, p < 0.001); Peak DR: (χ^2^(4) = 26.60, p < 0.001); TQ @ Drcrt: (χ^2^(4) = 64.94, p < 0.001); ADR: (χ^2^(4) = 136.06, p < 0.001); DR @ Drcrt: (χ^2^(4) = 5.04, p = 0.284).

